# Genetic canalization of nutrient resorption: evidence from a widespread grass under effective salt stress

**DOI:** 10.64898/2026.05.19.726328

**Authors:** Lele Liu, Wenyi Sheng, Huijia Song, Cui Wang, Na Li, Lele Lin, Yaolin Guo, Ning Du, Weihua Guo

## Abstract

Nutrient resorption is a key plant strategy for nutrient conservation, yet whether it responds plastically to non-nutrient stressors such as salinity remains unresolved. Using a common garden experiment with 110 genotypes of the cosmopolitan grass *Phragmites australis*, we first established that the imposed salinity treatment induced strong multi-level stress responses: above-ground biomass declined by >60%, sodium accumulated 3- to 5-fold across all leaf stages, and 484 metabolites showed significant differential accumulation, including canonical markers of osmotic and oxidative stress. Against this backdrop of confirmed stress, we found that nutrient resorption efficiency (NuRE) remained largely unaffected by salinity. Instead, NuRE was strongly correlated with phylogeographic lineage, ecotype, and latitude of origin, demonstrating evolutionary canalization rather than short-term acclimation. Element-specific regulatory patterns were also evident: while phosphorus resorption followed concentration-dependent control regardless of stress, nitrogen control was disrupted under salinity, and potassium resorption showed no such dependence. Our findings reveal that intraspecific variation in nutrient resorption is predominantly shaped by historical adaptation and geographic context, not by plasticity to salinity. This genetic canalization of a key functional trait implies that predictions of nutrient cycling under global change must account for the phylogeographic composition of plant populations.

## 1. Introduction

Nutrient resorption, the process by which plants translocate nutrients from senescing leaves to perennial tissues, is a pivotal conservation strategy that reduces dependence on soil nutrients and enhances nutrient use efficiency (Drenovsky et al., 2019; Jiang et al., 2023; Killingbeck, 1996). This mechanism is especially critical in nutrient-poor or stressful environments, influencing plant fitness, competitive ability, and ecosystem nutrient cycling (Yang et al., 2025). As a key functional trait, nutrient resorption efficiency (NuRE) reflects adaptive responses to environmental conditions. A suite of drivers, operating across environmental and plant-intrinsic levels, governs nutrient resorption. Environmentally, soil nutrient availability, temperature, and water stress are established abiotic modulators (Chen et al., 2021). More profoundly, plant-intrinsic control is conceptualized through several non-exclusive physiological hypotheses (Kobe et al., 2005; Sun et al., 2023): the nutrient concentration control posits that resorption efficiency declines as green leaf nutrient concentration rises; the stoichiometric control hypothesis suggests nutrients are resorbed proportionally to maintain internal balance; and the nutrient limitation control predicts plants preferentially resorb the nutrient that most limits their growth. Understanding how these mechanisms operate within species across diverse populations remains an important objective in functional ecology.

Intraspecific variation may arise from phenotypic plasticity in response to immediate stressors (Ramirez-Parada et al., 2024; Zhang et al., 2022), or from inherent genetic differences among populations shaped by evolutionary history and local adaptation (Ferrer Obiol et al., 2025; Radersma et al., 2020). Disentangling their contribution to NuRE is essential for predicting adaptive potential under environmental change (Zhang et al., 2022). However, most research on nutrient resorption has focused on its adaptation to nutrient limitation. Critically, its response to non-nutrient stresses, which disrupt plants through fundamentally different mechanisms like ionic toxicity and osmotic stress, remains a significant knowledge gap (Luo et al., 2021). Testing this gap is urgent because salinity stress is becoming a pervasive global change factor, albeit driven by distinct processes in different ecosystems (Zuo et al., 2024).

In coastal zones, salinity intrusion is primarily accelerated by sea-level rise and storm surges, threatening foundational wetland species and their biogeochemical functions (Seibert et al., 2024). In parallel, across arid and semi-arid inland regions, secondary soil salinization due to irrigation and groundwater over-exploitation is a major driver of land degradation and ecosystem alteration (Hassani et al., 2021). Despite these divergent drivers, the resultant physiological challenge to plants, ionic imbalance and water deficit, is convergent. This convergence provides a unique opportunity to investigate whether plants employ general or context-specific nutrient conservation strategies when facing salt stress, irrespective of its geographic origin.

Critically, predicting the direction of nutrient resorption responses to salinity remains challenging, rooted in two competing physiological and evolutionary perspectives (Luo et al., 2021; Zuo et al., 2024). On one hand, salinity imposes severe ionic and osmotic stress, disrupting membrane integrity, enzyme activity, and energy metabolism (Zuo et al., 2024). This could force a re-allocation of resources and potentially increase nutrient resorption efficiency as a compensatory conservation strategy to mitigate internal nutrient imbalance caused by uptake inhibition, consistent with the principles of nutrient limitation control (Kobe et al., 2005; Sun et al., 2023). Alternatively, nutrient resorption, particularly its stoichiometric relationships (e.g., N:P), may be under strong internal homeostasis control or subject to evolutionary canalization as a core nutrient conservation trait (Alseekh et al., 2025; Huang et al., 2022; Nicotra et al., 2010). Here, canalization denotes genetically based differences in NuRE among populations, coupled with limited phenotypic plasticity of this trait under the short-term salinity treatment. If this is the case, resorption patterns may exhibit conservatism, remaining unaltered by short-term abiotic stressors like salinity, with variation being primarily dictated by inherent genetic differences among populations shaped by long-term adaptation (Liu et al., 2025; Ren et al., 2020). Disentangling these possibilities (plastic acclimation versus inherent conservatism) is fundamental (Drenovsky et al., 2019; Prieto & Querejeta, 2020). It addresses whether this key functional trait (Killingbeck, 1996) can provide short-term flexibility to individual plants facing rapid environmental change, or whether ecosystem-level nutrient cycling responses will be constrained and predictable primarily from the genetic composition of plant populations.

These competing predictions can be tested by examining trait variation across natural geographic gradients. Geographic patterns, particularly across latitudinal and saline habitat gradients, offer a powerful lens to decipher the drivers of NuRE (Liu et al., 2025; Ren et al., 2020). In plant ecology, latitudinal clines in functional traits often reflect adaptive responses to overarching climatic filters such as temperature, growing season length, and nutrient availability (Li & Prentice, 2024). For nutrient resorption, broad-scale biogeographic studies have frequently documented increased efficiency at higher latitudes, a pattern interpreted as a conservative strategy to cope with shorter growing seasons and more constrained nutrient mineralization (Sun et al., 2016; Vergutz et al., 2012). However, in the context of salinization, an additional layer of complexity emerges: adaptive traits may be filtered not only by climate but also by historically saline conditions. Populations from coastal and inland salt marshes, despite their vast geographic separation, may have undergone convergent evolution for salt tolerance, potentially shaping similar nutrient management strategies. Disentangling whether observed trait variation aligns more strongly with broad-scale climatic gradients or with local adaptive pressures from saline habitats is therefore critical.

The common reed (*Phragmites australis*) is a uniquely powerful model to explore such intraspecific variation (Liu et al., 2025; Ren et al., 2020). It exhibits extensive geographic distribution and occupies the full spectrum of habitats relevant to this question, from freshwater wetlands to coastal saltmarshes and inland plateau saltmarshes, with distinct ecotypes and phylogenetic lineages demonstrating significant phenotypic divergence (Meyerson et al., 2025). By including populations from both coastal and inland saline origins, our experimental design explicitly captures the potential for convergent adaptation to salt stress, while also controlling for broad phylogeographic history. This allows us to separate the effects of evolutionary lineage, local habitat adaptation (salinity), and macro-climatic (latitudinal) gradients on a key functional trait.

In this study, we used a common garden experiment with multiple populations of *P. australis* spanning major phylogeographic groups and ecotypes (coastal vs. inland saline vs. freshwater) to: (1) first verify that the salinity treatment induces multi-level physiological stress (via biomass, Na⁺ accumulation, and metabolomics), a prerequisite for any meaningful genotype-by-environment analysis; (2) test whether the classic resorption control hypotheses are maintained or disrupted under salt stress; (3) quantify the relative contributions of phenotypic plasticity (to salinity) and population origin (phylogeography and ecotype) to NuRE variation, thereby directly assessing the ‘plastic acclimation versus inherent conservatism’ paradigm; and (4) decipher whether geographic variation in NuRE is primarily aligned with broad-scale climatic (latitudinal) gradients. We hypothesize that NuRE is a canalized trait largely unresponsive to short-term salt stress, and that its intraspecific variation is predominantly shaped by genetic origin and historical adaptation. By clarifying the mechanistic basis of intraspecific variation, this work seeks to scale from plant nutrient-use strategies to their potential impacts on ecosystem nutrient dynamics, improving predictions for ecological outcomes of salinization.

## 2. Materials and Methods

### 2.1 Common garden design

The common garden experiment was conducted in Changle County, Weifang City, Shandong Province, China (36°22′ N, 118°53′ E). On 5th April 2024, we collected 110 individuals (genotypes) of *P. australis* rhizomes from the common garden for cultivation. To disentangle the effects of deep evolutionary history from local adaptation, we employed a dual classification of our plant materials. The 110 individuals were assigned to six phylogeographic groups (AU, CN_N, CN_E, EU, EU_NA, NA; Table S1) based on genetic evidence from chloroplast haplotypes, nuclear microsatellites, and genomic analyses (Liu et al., 2022; Saltonstall, 2002; Wang et al., 2024). Concurrently, the populations from China were categorized into three ecotypes reflecting their source habitats: freshwater wetland, coastal saltmarsh, and inland plateau saltmarsh (Table S1). Ecotype classification was not applied to non-Chinese populations due to insufficient habitat documentation at the time of collection.

The plants were planted in barrels (total volume 25 L; top diameter 32.5 cm, bottom diameter 28.4 cm, height 38.5 cm) containing 20 L of a substrate composed of soil, peat moss, and river sand in a 2:1:1 volume ratio (Figure S1). The salinity treatment was applied as a 1% NaCl solution (10 ppt), calculated based on the estimated porewater volume of the 20 L soil system. Each individual was planted in two barrels: one for the salt treatment and one for the control. This concentration falls within the range encountered by *P. australis* in saline wetlands, and represents an ecologically recurrent stress that is moderate for most lineages of *P. australis* but severe for specific genotypes (e.g., those native to North America) (Eller et al., 2017; Meyerson et al., 2025), thus capturing the natural variation in salinity tolerance across its global distribution. This was achieved by adding 100 g of NaCl dissolved in water to each barrel on June 2 and 9, for a total of 200 g. Green leaves and senesced litter were sampled for elemental analysis at two critical phenological stages: one month after salt addition (July 11, 2024; coinciding with active growth) and six months later (December 4, 2024; coinciding with seasonal senescence), respectively.

At each sampling, we collected fully expanded, healthy green leaves from the top of the shoots, typically the 2nd to 4th leaf from the apex. Senesced litter was identified as leaves that had completely yellowed, which typically corresponded to the same 2nd to 4th positional nodes prior to senescence. This standardized protocol ensured the comparison of equivalent physiological tissues across all individuals and treatments.

### 2.2 Elemental and metabolomic profiling

The plant leaves and litter were dried at 45 ℃ to constant weight, cut with scissors, placed in a 2ml centrifuge tube and placed in two small steel balls, and ground with a grinder (Shanghai Jingxin Industrial Development Co., Ltd., Shanghai, China) at 55 Hz for 5 min to powder. Total carbon (C) and nitrogen (N) of leaves was measured using an element analyzer (UNICUBE, Germany) after weighing 2 mg (accurate to three decimal places) of plant samples in tin paper. A total of 0.1 g of the sample was weighed and placed in a polytetrafluoroethylene digestion tube, to which nitric acid was added, and then digested by microwave to obtain the digestive solution. The concentrations of K and P in the digest were determined by ICP-OES (iCAP Pro, Thermo Scientific, Germany).

Besides C, N, P and K, other seven elements (Cu, Zn, Fe, Mn, Mg, Si, Na) were quantified in leaf tissues. Concentrations of Na, Si, Fe, Mn, and Mg were determined by inductively coupled plasma optical emission spectrometry (ICP-OES; iCAP Pro, Thermo Scientific, Germany), while Cu and Zn were measured by inductively coupled plasma mass spectrometry (ICP-MS; NexION 1000G, PerkinElmer, USA).

For untargeted metabolomic profiling, approximately 50 mg of fresh leaf tissue was extracted with 1,000 μL of ice-cold extraction solvent (methanol:acetonitrile:water = 2:2:1, v/v/v) containing 20 mg/L of 2-chloro-L-phenylalanine as an internal standard. The mixture was homogenized with steel beads (45 Hz, 10 min), sonicated in an ice-water bath (10 min), and incubated at –20°C for 1 h. After centrifugation (12,000 rpm, 15 min, 4°C), the supernatant was vacuum-dried, and the residue was reconstituted in 160 μL of acetonitrile:water (1:1, v/v). Chromatographic separation was performed on an Acquity UPLC HSS T3 column (1.8 μm, 2.1 × 100 mm) using a Waters Acquity I-Class PLUS UHPLC system coupled to a Waters Xevo G2-XS QTOF mass spectrometer. The mobile phases consisted of 0.1% formic acid in water (A) and 0.1% formic acid in acetonitrile (B) at a flow rate of 400 μL/min with the following gradient: 0–0.5 min, 5% B; 0.5–5.5 min, 5–50% B; 5.5–9.0 min, 50–95% B; 9.0–10.5 min, 95% B; 10.5–12.0 min, 95–5% B. The injection volume was 2 μL. Mass spectrometry was operated in both positive and negative ion modes with an electrospray ionization source; capillary voltage was set to 2500 V (positive) or –2000 V (negative), cone voltage to 30 V, ion source temperature to 100°C, desolvation gas temperature to 500°C, cone gas flow to 50 L/h, and desolvation gas flow to 800 L/h. Data were acquired in MSe mode over a mass range of m/z 50–1200, alternating between low (off) and high (10–40 V) collision energies. Raw data were processed using Progenesis QI software for peak extraction, alignment, and annotation against the online METLIN database, public databases, and an in-house library.

For statistical analysis, raw metabolomic data were log₂-transformed, missing values were imputed with half of the minimum positive value, and constant metabolites were removed. Data were autoscaled (mean-centered and scaled to unit variance) prior to principal component analysis (PCA). Permutational multivariate analysis of variance (PERMANOVA) with 9,999 permutations was performed on Euclidean distances to test the effect of salinity while accounting for genotype as a covariate. Orthogonal partial least squares discriminant analysis (OPLS-DA) was used to extract variable importance in projection (VIP) scores. Differentially accumulated metabolites between salt and control groups were identified using limma with empirical Bayes moderation; significance was defined as |log₂ fold change| > 1, false discovery rate (FDR) < 0.05, and VIP > 1.

### 2.3 Nutrient resorption calculation

The continuous growth strategy and substantial leaf mass heterogeneity in *P. australis* preclude reliable mass loss correction factor (MLCF) determination. Uncontrolled physical losses during asynchronous senescence (notably a high rate of leaf tip abscission) decouple dry mass reduction from physiological nutrient resorption. Therefore, to ensure equitable comparisons, nutrient concentration in senesced leaves were standardized for carbon-basis shifts using the fractional change in the measurement basis (FCMB) protocol. Specifically, litter nutrient concentrations were multiplied by the ratio of green leaf carbon content to litter carbon content (Cgreen / Clitter) (Prieto & Querejeta, 2020). The Cgreen/Clitter ratios used for standardization varied among samples, with a mean of 1.06; a one-sample t-test confirmed this mean was significantly greater than 1 (*p* < 0.001), justifying the correction for a systematic shift in the carbon measurement basis. This approach directly quantifies nutrient withdrawal per structural carbon unit, neutralizing mass heterogeneity through carbon-standardization, bypassing fragmentation artifacts, and remaining valid across continuous growth gradients. Finally, nutrient resorption efficiency (NuRE) for N, P, and K was calculated as the percentage of nutrient recovered from senescing leaves: NuRE = (Green nutrient concentration - Litter nutrient concentration) / Green nutrient concentration × 100. Raw NuRE was calculated without carbon standardization and compared with carbon-standardized NuRE using Pearson correlations; the main linear mixed-effects analyses were also repeated using raw NuRE.

### 2.4 Data analysis

A principal component analysis (PCA) was conducted to visualise the overall structure of multi-element composition (N, P, K) across leaf types and salinity treatments, using the *rda* function in the *vegan* package after standardising the data.

Prior to multivariate analysis, all eleven elemental concentrations were standardized to zero mean and unit variance. A global principal component analysis (PCA) was performed using the *FactoMineR* package in R. To evaluate the overall effect of salinity on elemental composition, a multivariate analysis of variance (MANOVA) was conducted on PC1 and PC2 scores using Pillai’s trace test. Pairwise comparisons between control and salt treatments were performed using Welch’s *t*-test for PC1 and PC2 separately. To assess whether salinity effects differed across genetic lineages and developmental stages, MANOVA was repeated for each lineage × stage combination.

Phylogeographic group and ecotype were analysed in separate linear mixed-effects models because ecotype classifications were available only for Chinese populations and were partly associated with phylogeographic structure. Each model included salinity treatment and either phylogeographic group or ecotype as fixed effects, with genotype fitted as a random effect to account for the paired experimental design in which each genotype was exposed to both control and salt conditions. When significant effects were detected (*p* < 0.05), post-hoc pairwise comparisons were performed using Tukey’s HSD test via the *emmeans* package, with additional sensitivity analyses conducted on the within-genotype difference (Δ) between salt and control treatments.

The three fundamental control strategies over leaf nutrient resorption were assessed following established conceptual frameworks. To assess the nutrient concentration control strategy, we analyzed the relationships between log10-transformed nutrient concentrations in green leaves and senesced litter. A slope (*β*) significantly greater than 1 was interpreted as evidence of nutrient concentration control, indicating that resorption efficiency decreases as green leaf nutrient concentration increases. Conversely, a slope not significantly different from or less than 1 suggested the absence of this control strategy. To evaluate the stoichiometric control strategy, we examined the relationships between N and P resorption efficiencies, as well as between the resorbed N:P ratio and the N:P ratio in green leaves (Yang et al., 2025). Consistent positive correlations in both relationships were taken as support for stoichiometric control, reflecting proportional resorption aligned with initial nutrient ratios in green tissues. Non-significant or negative correlations indicated a deviation from this strategy. For the nutrient limitation control strategy, we tested the relationship between log10-transformed ratios of resorbed nutrients (e.g., N:P) and log10-transformed ratios in green leaves (Kobe et al., 2005). A slope significantly less than 1 was considered evidence of nutrient limitation control, implying that the resorption efficiency ratio between elements shifts in response to their relative limitation. A slope ≥ 1 suggested the ‘inverted’ nutrient limitation control (Sun et al., 2023).

To rigorously estimate the functional (structural) slopes between bivariate nutrient traits (e.g., green leaf vs. senesced litter concentrations), we employed Standardized Major Axis (SMA) regression implemented via the *smatr* package in R (Warton et al., 2012). SMA regression was selected as the primary analytical method because it is a Model II regression approach that appropriately accounts for measurement and biological variation in both variables—a critical consideration given the observational nature of our data where both nutrient concentrations in green leaves and senesced litter are measured with error. This method provides unbiased estimates of the true structural relationship between variables, unlike Ordinary Least Squares (OLS) regression which assumes the predictor variable is error-free and systematically underestimates slope magnitudes under these conditions (Warton et al., 2012).

For each bivariate relationship, we performed SMA regression analyses separately for each salinity level. The *smatr* package was used to estimate the SMA slope (*β*) along with its 95% confidence interval and to calculate the coefficient of determination (*R*²). Crucially, we specifically tested whether each estimated SMA slope significantly deviated from theoretical values, particularly from 1. This slope comparison against 1 was conducted using the built-in permutation tests in the *smatr* package, which generated *p*-values indicating the statistical significance of the deviation. For comparative purposes and to maintain consistency with the methodological approaches employed in foundational studies of nutrient resorption (e.g., Kobe et al., 2005; Sun et al., 2023) that relied on OLS regression, we also calculated and report OLS slope estimates in supplementary materials. However, all formal hypothesis testing and interpretation regarding slope values, particularly tests of deviation from 1:1 relationships, are based on the more appropriate SMA regression results, which represent robust, unbiased estimates of the functional relationships between nutrient traits.

The influence of geographical origin (absolute latitude) on NuRE was analysed using linear mixed-effects models (LMMs) fitted with the *lmer* function in R package *lme4* (Bates et al., 2015), where phylogeographic group was included as a random intercept term to account for non-independence among individuals from the same source. Model assumptions of normality and homoscedasticity were verified by inspecting residual plots, and NuRE values were arcsine-square-root transformed when necessary to meet these assumptions. To quantify the individual contributions of the fixed factors (latitude, leaf nutrient concentration, and stoichiometry) to the variance in NuRE, we performed a variation partitioning analysis flowing the LMMs with phylogeographic group as a random intercept. This variation partitioning was conducted using the *glmm.hp* package in R (Lai et al., 2022), which implements hierarchical partitioning to calculate the marginal *R*^2^ for each predictor within the fitted linear mixed-effects model framework. This approach allowed us to estimate the unique proportion of variance in NuRE attributable to each fixed factor, independent of the random effect of phylogeographic group. All figures were generated using ggplot2 and assembled with the patchwork library. All statistical analyses were performed in R version 4.3.0.

## 3. Results

### 3.1 Salt stress induces multi-level physiological responses

To establish that the salinity treatment imposed a physiologically meaningful stress before evaluating NuRE responses, we first examined multi-level indicators of salt stress

Principal component analysis (PCA) of the metabolome revealed a clear separation between salt-treated and control plants (Figure 1a), with the first two principal components explaining 16.51% and 7.81% of the total variance, respectively; orthogonal partial least squares discriminant analysis yielded a *Q*²(cum) of 0.233, indicating that the treatment effect is detectable albeit accompanied by substantial genotypic variation. A total of 484 metabolites showed significant differential accumulation between treatments (|log₂ fold change| > 1, *FDR* < 0.05, *VIP* > 1) (Figure 1b). Among the metabolites, canonical stress markers were prominently altered: N,N-dimethylglycine (a glycine betaine precursor) and sucrose 6-phosphate were significantly upregulated (log₂FC = +1.04 and +2.93, respectively), while glutathione (log₂FC = –0.27) and N-caffeoylputrescine (log₂FC = –1.34) were significantly downregulated. Collectively, these results demonstrate that the salinity treatment induced widespread metabolic reprogramming consistent with osmotic and oxidative stress responses.

**Figure 1.**
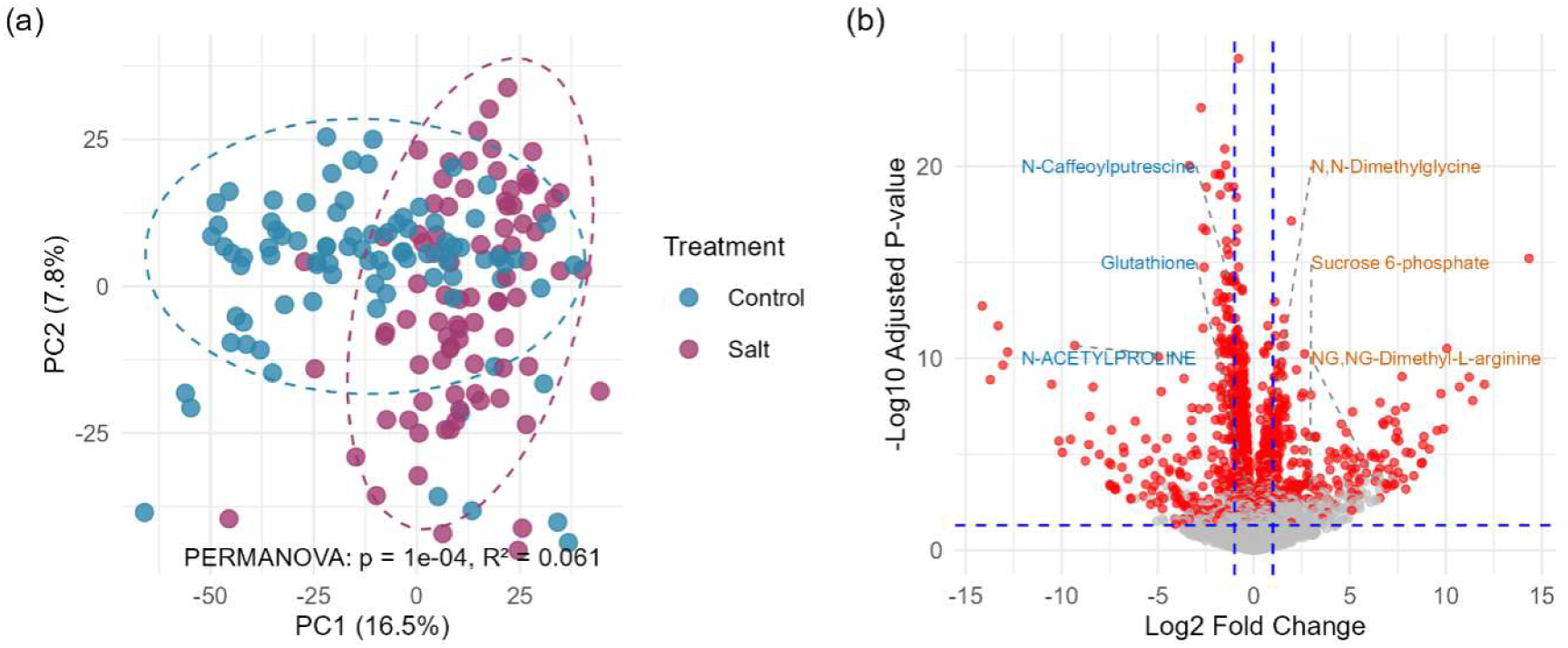
Metabolomic responses of *Phragmites australis* to salinity stress. (a) Principal component analysis (PCA) score plot of the metabolome (3,413 metabolites) under control (blue) and salt (red) treatments. Ellipses represent 95% confidence intervals for each group. PERMANOVA (Euclidean distance, 9,999 permutations) with genotype as covariate revealed a significant effect of salinity (p < 0.001). (b) Volcano plot showing differential accumulation of metabolites between salt and control groups. Red points indicate metabolites that are significantly different (|log₂ fold change| > 1, false discovery rate < 0.05, variable importance in projection > 1). Selected stress-responsive metabolites (e.g., N,N-dimethylglycine, sucrose 6-phosphate, glutathione) are labeled.

Salt treatment drastically reduced above-ground biomass by more than 60% (control: 835.7 ± 36.4 g, salt: 310.4 ± 15.0 g; Welch’s *t* = 13.26, *p* < 0.001; Figure 2a). Sodium (Na) accumulation, a direct indicator of salt uptake, increased dramatically across all leaf developmental stages. In green leaves, Na concentrations rose from 475 to 1929 mg kg⁻¹ *p* < 0.001), in yellowing leaves from 508 to 2027 mg kg⁻¹ (*p* < 0.001), and in litter leaves from 1379 to 3722 mg kg⁻¹ (*p* < 0.001; Figure 2b).

**Figure 2.**
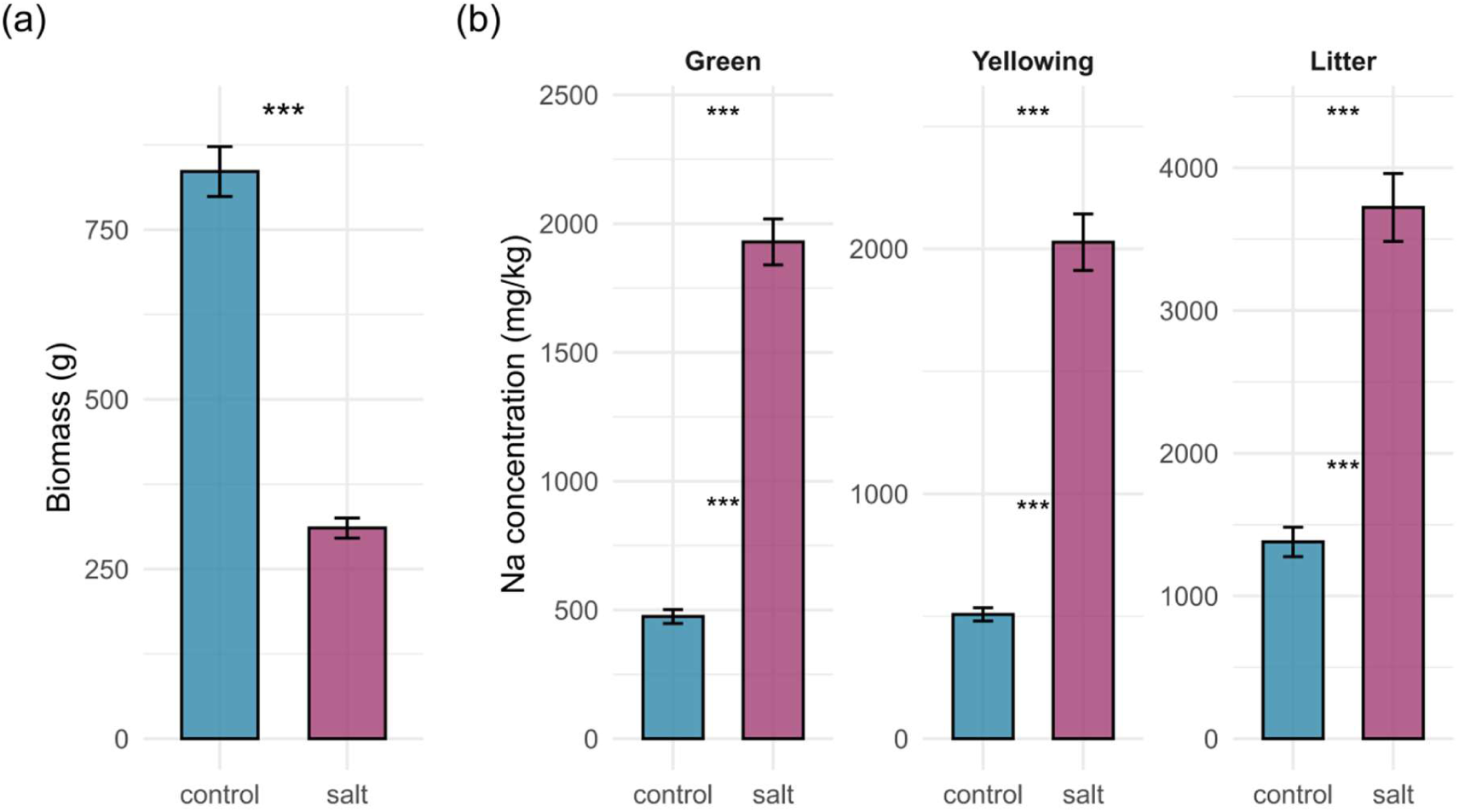
Effects of salinity on biomass and sodium accumulation in *Phragmites australis*. (a) Biomass of control and salt-treated plants (harvest at the final stage). Data are means ± SE. *** *p* < 0.001 (Welch’s *t*-test, *n* = 106 per group). (b) Sodium (Na) concentrations in green, yellow, and brown leaves under control and salt conditions. Data are means ± SE. *** *p* < 0.001 (Welch’s t-test for each stage, *n* = 105–106 per group per stage).

The elemental composition of leaves was also significantly altered by salinity. Principal component analysis of 11 elements revealed that PC1 (41.9% variance) separated nutrient–related elements (C, N, P, K, Cu, Zn) from stress–related elements (Si, Mn, Na), while PC2 (14.3%) was primarily driven by Mn, Zn, Cu, Fe, and Si (Figure S2a). Salinity significantly increased PC2 scores (*p* < 0.001, MANOVA), indicating a shift in elemental balance (Figure S2bc). Notably, the effect of salinity on elemental composition was stage– and lineage–dependent: during the Green stage, all lineages showed significant responses; during the Yellowing stage, only CN_N and EU lineages were affected; and by the Litter stage, no significant salinity effect was detected in any lineage (Figure S2d-f).

### 3.2 Nutrient resorption efficiency is genetically canalized under effective salt stress

The PCA of leaf N, P, and K concentrations showed a clear separation between green leaves and senesced litter along the first principal component (PC1, Figure S3a). This separation was driven by the strong positive loadings of all three nutrient concentrations on PC1 (Figure S3b). This pattern indicates that the dominant trend in the data is the distinction between nutrient-rich green leaves and nutrient-poor senesced litter, a pattern that directly results from substantial nutrient resorption during senescence.

The results of the linear mixed–effects models indicated a significant effect of phylogeographic group on the resorption efficiency of N, P, and K (*p* < 0.001); however, neither salinity nor the interaction between salinity and phylogeographic group had a significant effect (Figure 3; Figure S4). For instance, the CN_N group exhibited the lowest N and P resorption efficiencies among all groups, whereas both the CN_N and CN_E groups showed high K resorption efficiency. Meanwhile, the results revealed a significant effect of ecotype on the resorption efficiency of N and P (*p* < 0.001), but not on K (*p* = 0.694) (Figure 4; Figure S5). It should be noted that the ecotype analysis here was based only on Chinese populations, as ecotype classification was not available for non-Chinese populations. A significant interaction between salinity and ecotype was observed for N resorption efficiency (*p* = 0.038), indicating that salinity treatment enhanced N resorption in the saltmarsh ecotype relative to the freshwater ecotype. Among the ecotypes, coastal saltmarsh populations displayed the highest N and P resorption efficiencies, while populations from inland plateau saltmarsh had the lowest.

**Figure 3.**
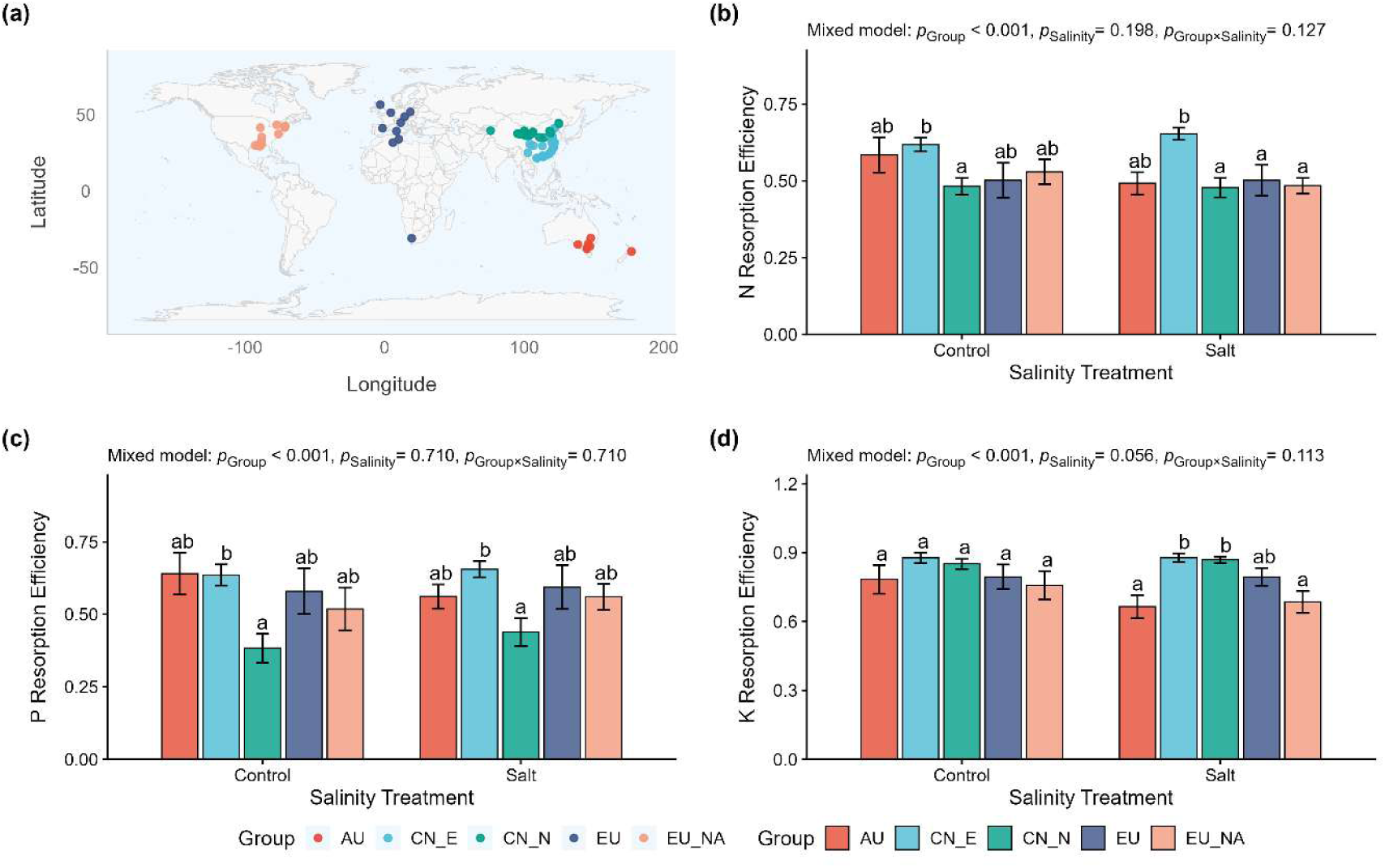
Geographical distribution (a) of the phylogeographic groups of *Phragmites australis* and leaf nutrient resorption efficiency (b–d) for nitrogen (N), phosphorus (P), and potassium (K). Phylogeographic groups reflect deep evolutionary lineages defined by genetic markers (e.g., chloroplast haplotypes). Statistical significance was assessed using linear mixed-effects models that included genotype as a random effect to accommodate the paired experimental design. The *p*-values shown in panels b–d refer to the fixed effects of group, salinity and their interaction. When significant group effects were detected, post-hoc pairwise comparisons were performed separately within each salinity treatment using Tukey’s HSD test; different letters denote statistically significant differences (*α* = 0.05).

**Figure 4.**
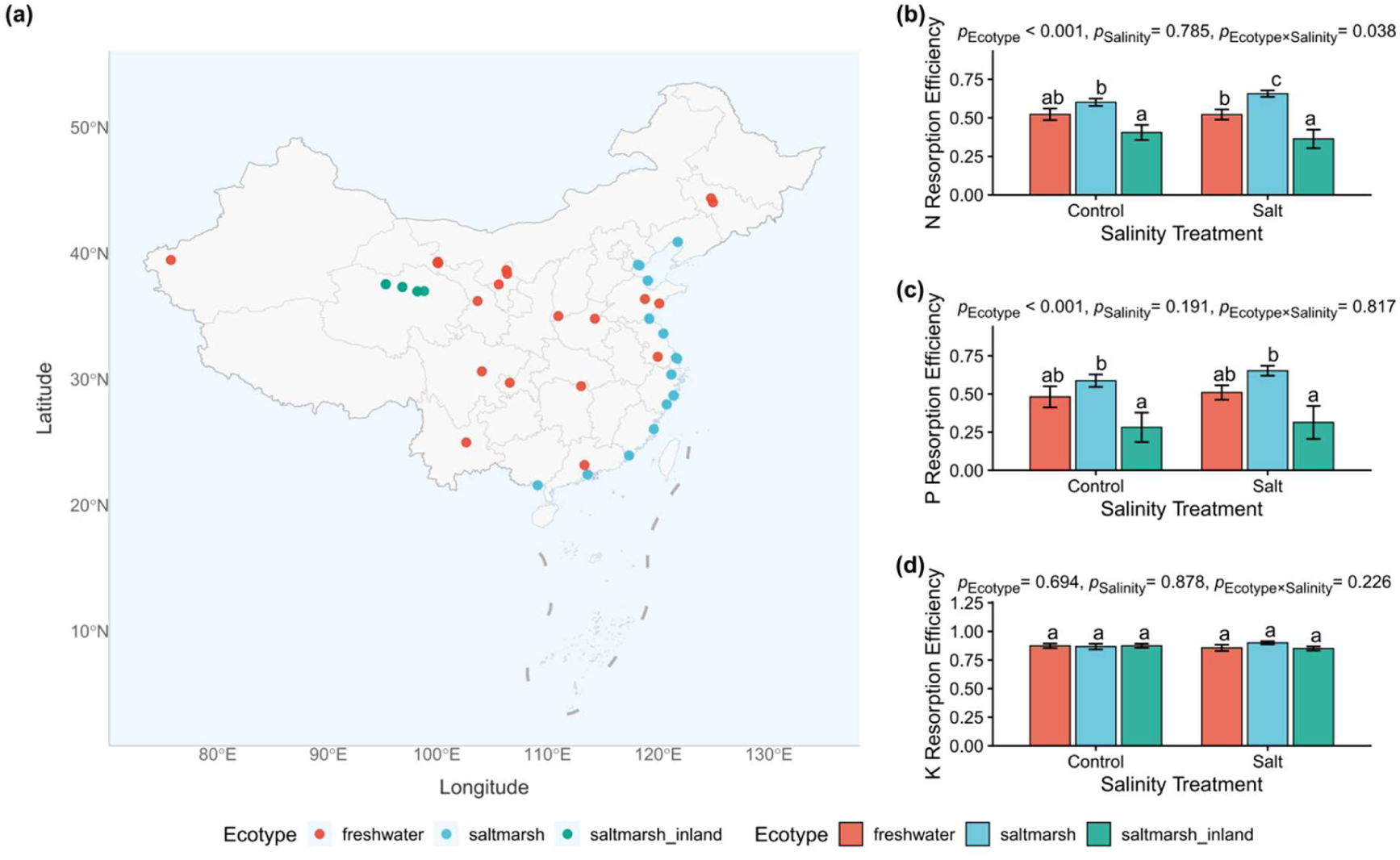
Geographical distribution (a) of *Phragmites australis* ecotypes and leaf nutrient resorption efficiency (b–d) for nitrogen (N), phosphorus (P), and potassium (K). The colour scheme represents habitat-based adaptations and is independent of that used in Figure 3. Ecotypes categorize populations according to adaptive phenotypes to specific local habitats (freshwater wetlands, coastal saltmarsh, inland saltmarsh). Significance was tested with linear mixed-effects models incorporating genotype as a random effect to account for paired measurements. The reported *p*-values correspond to the fixed effects of ecotype, salinity and their interaction. Where ecotype effects were significant, post-hoc Tukey’s HSD comparisons were conducted within each salinity level; different letters indicate statistically significant differences (*α* = 0.05).

### 3.3 Element-specific regulatory strategies govern nutrient resorption

A significant positive association was observed between N concentrations in green leaves and litter under control conditions (*β* > 1, *p* < 0.05; Figure 5a), indicating the presence of nutrient concentration control over leaf N resorption in *P. australis* under non-saline conditions. However, this relationship disappeared under salt treatment (*p* > 0.05; Figure 5a), suggesting that salinity disrupted the N concentration control strategy. For P, a significant positive association was maintained under both control and salt conditions (*β* > 1, *p* < 0.05; Figure 5b), indicating robust nutrient concentration control over leaf P resorption in *P. australis* regardless of salinity. In contrast, correlations for K were non-significant under both conditions (*p* > 0.05; Figure 5c), suggesting the absence of nutrient concentration control for this element.

**Figure 5.**
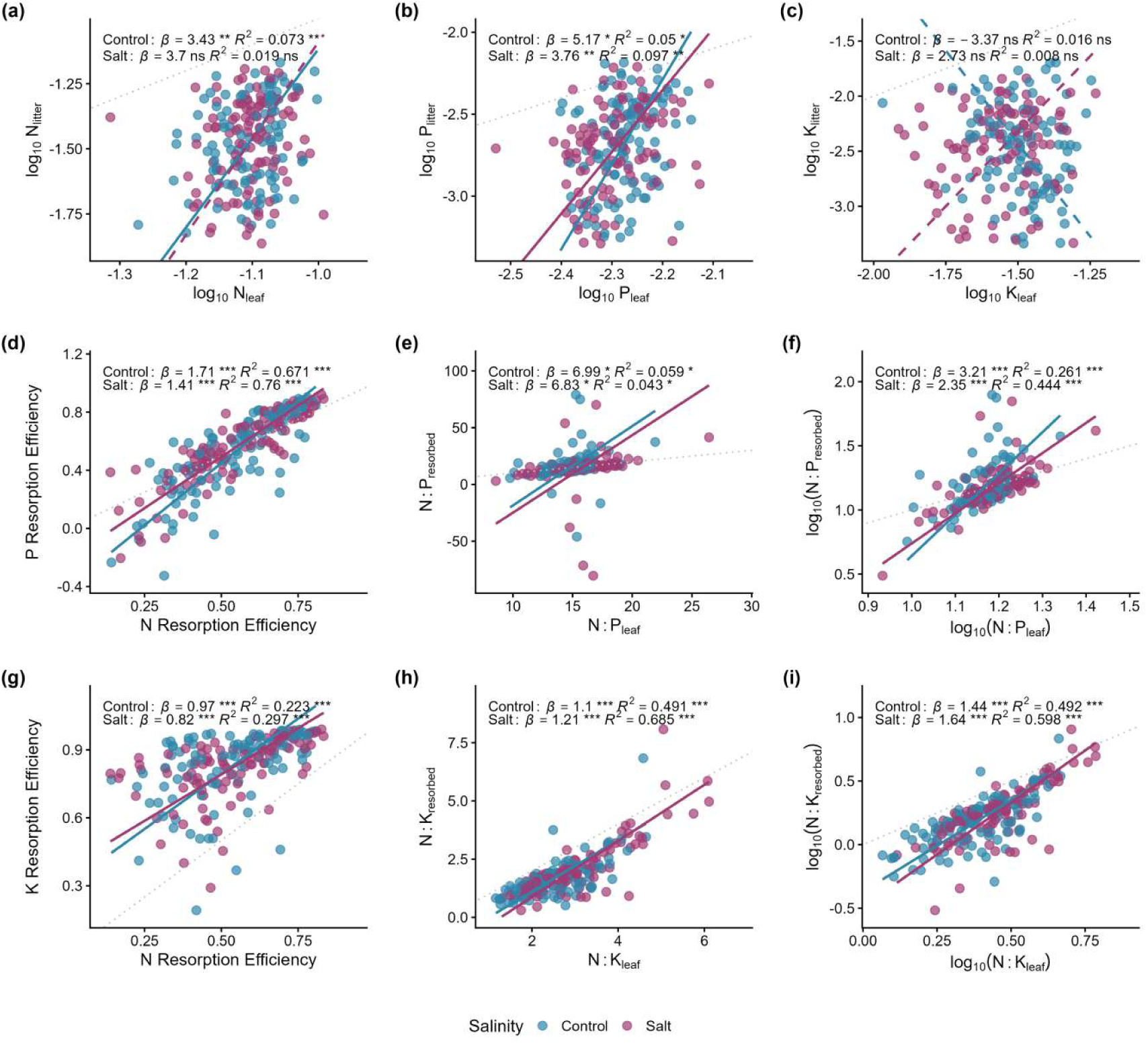
Assessment of three control strategies for nutrient resorption in *Phragmites australis* under varying salinity levels. (a–c) Relationship between nutrient concentrations (N, P, K) in green leaves and senesced litter. (d, g) Association between phosphorus and nitrogen resorption efficiencies. (e, h) Linkage between resorbed N:P(K) ratios and green leaf N:P(K) ratios. (f, i) Correlation between log₁₀-transformed resorbed N:P(K) and log₁₀-transformed green leaf N:P(K) ratios. The gray dashed line in each panel indicates the 1:1 reference. Solid lines represent significant relationships (*p* < 0.05 for the SMA regression model), while dashed lines indicate non-significant relationships. Statistical annotations show results from standardized major axis (SMA) regression for each salinity level: *β* (with asterisks) is the SMA slope, where asterisks after *β* indicate significant deviation from 1 (* *p* < 0.05, ** *p* < 0.01, *** *p* < 0.001), and asterisks after *R*² indicate significance of the SMA regression model.

Significant positive relationships were detected between N and P resorption efficiencies (Figure 5d), as well as between the resorbed N:P ratio and the green leaf N:P ratio under both control and salt conditions (Figure 5e), supporting the presence of stoichiometric control over P resorption. Similarly, N and K resorption efficiencies were positively correlated (Figure 5g), and the resorbed N:K ratio was significantly related to the green leaf N:K ratio under both conditions (Figure 5h), indicating stoichiometric control also regulates K resorption.

Additionally, significant linear relationships were found between the log₁₀-transformed resorbed ratio and log₁₀-transformed green leaf ratio for N:P and N:K, with slopes greater than 1 (Figure 5f, i), demonstrating an ‘inverted’ nutrient limitation control for P and K relative to N. The OLS slope estimates (Figure S6) generally aligned with these resorption control strategies inferred from the SMA regressions.

### 3.4 Latitudinal variation reflects macro-scale adaptation

The linear mixed-effects models revealed that NuREs of N, P, and K were significantly positively correlated with the absolute latitude of population origin (Figure 6). Variation partitioning indicated that latitude had the largest individual contribution among the measured predictors, but its contribution was modest for N, P, and K resorption (individual *R*² = 0.092, 0.057, and 0.094, respectively; Table 1). The very small contributions of green-leaf P concentration (*R*² = 0.006) and stoichiometry (*R*² < 0.001) further indicate that most variation in P resorption was associated with factors not represented in the present models.

**Figure 6.**
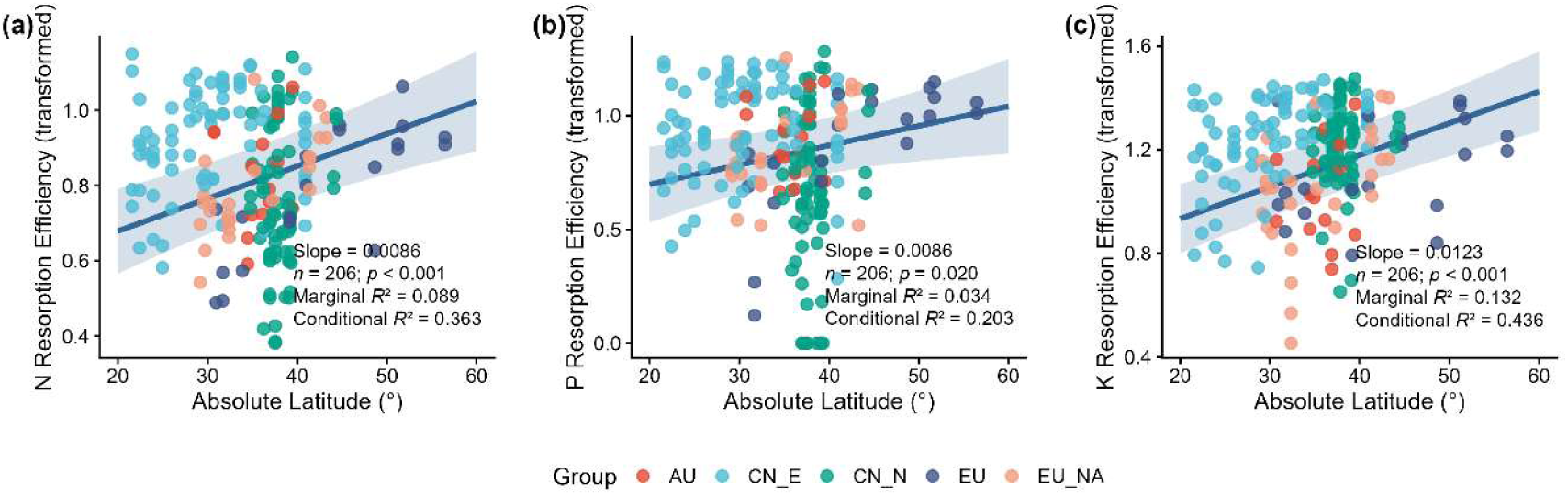
Influence of latitude on leaf nitrogen (N), phosphorus (P), and potassium (K) resorption efficiencies of *Phragmites australis*. The analysis was performed using linear mixed-effects models, treating phylogeographic groups as a random effect.

**Table 1.**
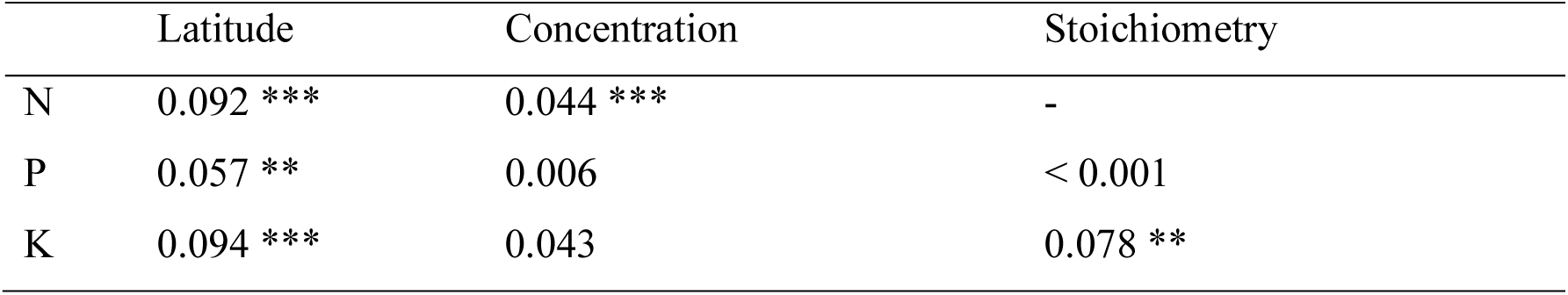
Individual effects (*R*²) of absolute latitude of origin, nutrient concentration, and nutrient stoichiometry on leaf N, P, and K resorption, estimated using linear mixed-effects models with phylogeographic group as a random intercept. Individual marginal *R*² values were calculated by hierarchical partitioning using *glmm.hp*. Asterisks indicate the significance of the corresponding fixed effects in the full models: * *p* < 0.05; ** *p* < 0.01; *** *p* < 0.001.

Raw and carbon-standardized NuRE were strongly correlated for N, P, and K (Pearson’s *r* = 0.997–0.999; Figure S7), and analyses using raw NuRE did not alter the results for phylogeographic group, ecotype, salinity, their interactions, or latitude.

## 4. Discussion

### 4.1 Effective salt stress meets conservative NuRE: reconciling the paradox

We first confirmed that the salinity treatment imposed a strong and sustained physiological stress at multiple biological levels. Metabolomic profiling revealed clear separation between control and salt-treated plants, with 484 metabolites differentially accumulated, including canonical markers of osmotic and oxidative stress such as N,N-dimethylglycine (a glycine betaine precursor) and sucrose 6-phosphate (Figure 1). Above-ground biomass declined by more than 60% (Figure 2a), and leaf Na concentrations increased 3- to 5-fold across all developmental stages (Figure 2b). Elemental composition also shifted significantly, with salinity increasing PC2 scores (mainly driven by Mn, Zn, Cu, Fe, Si) in a stage- and lineage-dependent manner (Figure S2). Collectively, these results confirm that the treatment was physiologically effective, providing a robust foundation to evaluate NuRE responses. In this study, metabolomic profiling was used primarily to confirm that the salinity treatment induced broad physiological stress, rather than to resolve genotype-specific metabolic mechanisms underlying NuRE variation. Integrating metabolite profiles with genotype-level NuRE responses would be a valuable direction for future mechanistic research. Against this background of confirmed stress, the intraspecific variation in nutrient resorption efficiency (NuRE) we observed in *P. australis* reflects divergent genetic origins, not phenotypic plasticity. Our results demonstrate that phylogeographic group and ecotype are primary determinants of NuRE, while salt stress elicited no overall plastic response in NuRE. This finding highlights the conservative nature of NuRE under this non-nutrient stress condition.

Nutrient resorption is established as a key plant functional trait governing nutrient conservation strategies and ecosystem processes (Killingbeck, 1996; Kobe et al., 2005). Therefore, deciphering the drivers of its intraspecific variation (genetic adaptation versus plasticity) is central to functional ecology (Liu et al., 2025; Zhang et al., 2022). Our finding that this variation is shaped by long-term adaptive evolution rather than a general short-term acclimation capacity underscores that macroecological patterns can arise primarily from genetic differentiation. Such trait conservatism, also observed in desert plants under saline stress (Luo et al., 2021), suggests evolutionary optimization under consistent selective pressures, which may constrain phenotypic adjustments to sudden environmental shifts. The predominance of genetic over plastic controls highlights the necessity of incorporating intraspecific diversity into future models of nutrient cycling.

The lack of a plastic increase in NuRE under salinity presents a physiological puzzle. Theoretically, if salt stress inhibits root nutrient uptake through ionic competition or root damage, enhancing resorption from senescing leaves could serve as a critical internal compensation mechanism to maintain minimum nutrient levels for growth and metabolism. This aligns with ecological principles of optimal allocation and stress adaptation, where optimizing internal recycling is a predicted strategy under resource limitation (Nicotra et al., 2010). However, our data did not support this expected plastic increase, a phenomenon also noted in some desert plants under saline stress (Luo et al., 2021). This discrepancy suggests the existence of substantive physiological constraints that override this optimal strategy. Salt stress may disrupt the finely tuned cellular senescence program necessary for efficient nutrient remobilization, possibly through the accumulation of incompatible solutes or oxidative damage that affect element stability (Zuo et al., 2024), or through a preferential reallocation of energy towards immediate survival mechanisms (e.g., osmotic adjustment and ion exclusion). Therefore, the canalization of NuRE may reflect an evolutionary ‘optimum’ under stress (Huang et al., 2022), where the metabolic and logistical costs of enhancing resorption outweigh its potential benefit, or where the core resorption machinery is intrinsically vulnerable to ionic interference (Alseekh et al., 2025).

Our findings appear to contrast with studies investigating other abiotic stressors. For example, a five-year field experiment simulating climate change, specifically warming and reduced rainfall, reported significant declines in resorption efficiency for multiple nutrients across several shrub species (Prieto & Querejeta, 2020). This discrepancy may stem from differences in the type and duration of stress applied. Our inference of canalization is therefore limited to the absence of a plastic NuRE response during one growing season under the imposed salinity treatment. Chronic, more severe, or multigenerational salinity exposure may produce acclimatory, epigenetic, or transgenerational responses that cannot be evaluated here. Moreover, although altered phloem loading, disruption of senescence-associated remobilization, and reallocation towards osmotic adjustment are plausible explanations for the observed response, we did not directly measure these mechanisms. Long-term experiments combined with targeted molecular and transport measurements are needed to distinguish among these possibilities. Because only one moderate salinity level (10 ppt) was tested, our results do not exclude plastic NuRE responses at higher salinity or along a dose-response gradient. Future experiments should determine whether a threshold exists beyond which the apparent stability of NuRE is overcome.

Alternatively, nutrient resorption may exhibit stressor-specific plasticity, being more responsive to soil nutrient availability and climate-related shifts than to ionic stress such as salinity (Drenovsky et al., 2019; Luo et al., 2021; Yang et al., 2025; Yuan & Chen, 2015). Moreover, the contrast underscores the potential importance of experimental duration (long-term gradual stress versus short-term shock) in shaping plant physiological responses. Such comparisons highlight the need for integrative approaches across stress types and time scales to fully understand the plasticity and evolution of nutrient conservation strategies.

### 4.2 Divergent strategies control the resorption of N, P, and K

Our study revealed distinct regulatory mechanisms governing the resorption of N, P, and K in *P. australis*. Phosphorus resorption was consistently influenced by initial green tissue P concentration under both control and saline conditions, supporting the nutrient concentration control hypothesis. In contrast, N resorption followed concentration control only under non-saline conditions, with this relationship disappearing under salinity. Potassium resorption showed no concentration dependence. Notably, N and K resorption exhibited tight stoichiometric coordination, as reflected in stable N:P and N:K ratios across treatments.

Functionally, slopes greater than 1 indicate that changes in green-leaf N:P or N:K are accompanied by disproportionate changes in the corresponding resorbed ratio, consistent with relatively greater recovery of P or K than of N across the observed nutrient gradient. This pattern may contribute to maintaining internal N:P:K balance during regrowth. It may also reflect stronger conservation of P, the high mobility of K, or constraints on the remobilization of N retained in structural or metabolic compounds. Because the experiment did not include nutrient additions or direct measurements of remobilization costs, these explanations remain hypotheses.

These element-specific strategies align with broader patterns documented across biomes and taxa. For instance, the prevalence of stoichiometric control for multiple nutrients, including N and K, has been demonstrated at a global scale (Sun et al., 2023), suggesting a universal mechanism that complements nutrient limitation effects. Notably, K resorption did not follow a concentration-dependent pattern, despite this element typically exhibiting high resorption efficiencies globally (often >70%; Vergutz et al., 2012). This suggests that high K recovery may be driven by factors other than green leaf K concentration. One possible explanation is the partial functional substitution of K⁺ by Na⁺ during osmotic adjustment. Na⁺ can replace part of the nonspecific vacuolar osmotic function of K⁺ in plants and has been reported to become a major osmoticum in *P. australis* from higher-salinity habitats (Wakeel et al., 2011; Zhao et al., 1999). Increased Na⁺ accumulation may therefore reduce reliance on K⁺ for osmotic adjustment, potentially contributing to the weak relationship between green-leaf K concentration and K resorption efficiency observed here. Furthermore, the preferential resorption of P under certain conditions echoes patterns found in nutrient-limited systems such as tropical forests and alpine permafrost ecosystems, where P resorption can surpass N resorption due to strong local constraints (Yang et al., 2025). This suggests that the strong concentration-dependent control of P resorption we observed may be a widespread adaptation to P limitation, which can override other biogeographic expectations. The stability of these stoichiometric relationships under salt stress implies that internal nutrient regulation is a conservative trait, potentially shaped by evolutionary history more than immediate abiotic context. Such stoichiometric flexibility may be especially critical in environments subject to periodic nutrient imbalances, enabling plants to maintain functional homeostasis.

It is important to acknowledge that our experimental design did not include a nutrient addition treatment, which represents the most direct approach to test the nutrient limitation control hypothesis (Chen et al., 2021). Future studies incorporating nutrient manipulations would provide more definitive evidence. However, our use of resorbed nutrient ratios (e.g., Resorbed N:P) as a proxy for relative nutrient limitation is a well-established methodology in resorption ecology (Kobe et al., 2005; Sun et al., 2023), and has been effectively applied in other recent studies (e.g., Yang et al., 2025). The significant intraspecific variation across our studied populations inherently created a broad gradient of nutrient status and demand. Within this context, the consistent ‘inverted’ nutrient limitation pattern (i.e., a slope significantly >1 for the relationship between log Resorbed N:P and log Green N:P) provides indirect evidence consistent with nutrient-limitation control at the intraspecific level in *P. australis*, but it cannot establish causal nutrient limitation. Direct factorial nutrient-addition experiments would be required to determine whether the observed resorption patterns are driven by the relative limitation of N, P, or K.

### 4.3 Latitudinal gradient in resorption reflects macro-Scale adaptation

Latitude consistently predicted nutrient resorption efficiency (NuRE) in our study, with populations from higher latitudes exhibiting greater resorption for all nutrients. This macroecological pattern likely reflects adaptive responses to environmental constraints such as shorter growing seasons, lower temperatures, and reduced nutrient availability, which collectively favor more conservative nutrient use strategies. Although latitude explained more variation than the other measured predictors, its individual contribution remained modest. The substantial unexplained variance indicates that additional climatic, edaphic, demographic, or genetic factors also contribute to NuRE variation. We therefore interpret latitude as a significant but limited correlate of NuRE rather than as a dominant determinant.

This gradient aligns with biogeographic patterns observed in other widely distributed species. For instance, increasing nutrient resorption efficiency with latitude has been documented in temperate trees such as *Quercus variabilis* across China, where N, P, K, and S resorption all increased significantly with latitude, paralleled by declining nutrient concentrations in senesced leaves (Sun et al., 2016). Such convergent evolution of nutrient conservation strategies across distinct lineages and biomes suggests a general principle in plant adaptation to high-latitude environments. However, the relative contribution of genetic differentiation versus phenotypic plasticity to these latitudinal clines may vary by species and ecosystem. For example, in the grass *Stipa breviflora*, phenotypic plasticity to soil nutrient availability was the dominant factor explaining latitudinal variation in resorption, with genetic effects playing a modest role (Zhang et al., 2022). This contrasts with our findings in *P. australis*, wherein geographic origin (reflecting genetic or deep-time adaptive processes) outweighed plastic responses to salinity. Therefore, while latitudinal clines in resorption traits are common, our study demonstrates that they can arise predominantly from genetic differentiation rather than phenotypic plasticity.

### 4.4 Ecological and evolutionary implications of genetic canalization

Our findings demonstrate that NuRE in *P. australis* is a genetically conservative trait with limited phenotypic plasticity to salt stress. This conclusion is reinforced by the clear evidence that the salinity treatment was effective at multiple levels (metabolic reprogramming, biomass reduction, sodium accumulation, and altered elemental balance) yet NuRE remained unchanged. Such a decoupling of a core functional trait from an effective environmental stress challenges the prevailing assumption of broad trait plasticity to environmental change (Nicotra et al., 2010) and underscores the role of evolutionary constraints, such as stabilizing selection, in canalizing this trait (Luo et al., 2021). Crucially, we show that while salinity can disrupt specific regulatory patterns (e.g., concentration control for N), the overall efficiency of nutrient recovery remains fixed and is predominantly governed by geographic origin (phylogeographic group and ecotype). The separate-model approach avoids including the two correlated classification schemes as simultaneous independent predictors, but it does not fully disentangle deep phylogeographic history from recent habitat-associated differentiation. We therefore interpret the ecotype analysis as complementary evidence of habitat-associated differentiation rather than as an effect independent of phylogeographic history.

This conservatism reframes how we predict ecosystem function under global change. Since trait variation is phylogenetically anchored, the pace and direction of ecosystem nutrient cycling changes will be forecast primarily by the pre-existing genetic composition of plant populations. In rapidly salinizing habitats, adaptive potential therefore relies on standing genetic variation rather than short-term acclimation, making the conservation of intraspecific genetic diversity critical for ecosystem resilience (Liu et al., 2021). To accurately project biogeochemical feedbacks, models must integrate intraspecific variation and phylogeographic history alongside environmental drivers (Vahsen et al., 2023).

These principles have direct management and biosecurity relevance. *P. australis* is widely introduced for wetland restoration and can be invasive (Meyerson et al., 2025). Our study shows that key functional traits like NuRE are strongly conserved within lineages. For instance, the invasive European lineage in North America (EU_NA) and its native progenitor (EU) showed no significant divergence in NuRE (Figure 3), a functional stasis consistent with observations of genetic and epigenetic stability during invasion (Liu et al., 2018). Consequently, the transfer or expansion of distinct genotypes (e.g., high-NuRE types) could establish populations with inherent, non-plastic nutrient cycling signatures in new ecosystems. Therefore, the functional traits of source populations must be a key criterion in restoration projects, and monitoring of invasive genotypes should account for their potential to alter local nutrient dynamics.

By evaluating multiple nutrients across diverse genetic lineages, our study reveals that classical resorption hypotheses operate as conditional and hierarchical regulatory strategies (Alseekh et al., 2025; Chen et al., 2021). Their influence is element-specific: concentration control firmly governs P resorption (Kobe et al., 2005) but is disrupted for N under salinity, while stoichiometric control persists throughout (Sun et al., 2023). This integrative framework moves beyond single hypotheses and offers a more realistic basis for predicting plant responses to complex stress. To advance this framework, future studies should: (a) combine nutrient additions with salt stress to test interactive effects on resorption control; (b) apply transcriptomic or metabolomic tools to elucidate the regulatory pathways underlying these canalized traits (Huang et al., 2022); and (c) examine whether conservative resorption is common observed across other foundation species. These steps are vital for assessing the adaptive capacity of plant communities in a changing world.

## 5. Conclusions

This study demonstrates that intraspecific variation in nutrient resorption efficiency in *P. australis* is primarily determined by genetic origin and geographic context, particularly latitude, rather than by plastic responses to non-nutrient stressors. Despite clear evidence of effective salt stress, including substantial biomass reduction, pronounced sodium accumulation, and widespread metabolic reprogramming, NuRE showed no overall plasticity. Instead, we identified distinct, element-specific control strategies: under control conditions, N resorption followed concentration control, a pattern disrupted by salt stress; P resorption was consistently regulated by initial tissue concentration across treatments; and both N and K resorption were under strong stoichiometric control, with evidence of inverted nutrient limitation for P and K relative to N. These patterns reflect evolutionary adaptations to local nutrient conditions and are likely conserved across widely distributed plant species. The limited phenotypic plasticity of NuRE under salt stress suggests that ecosystem resilience to rapid non-nutrient environmental change may depend more on standing genetic variation than on short-term acclimation. Consequently, integrating phylogenetic and macroecological perspectives, particularly the phylogeographic composition of plant populations, is essential for accurately forecasting future nutrient cycling dynamics in a changing world.

## Acknowledgements

This work is supported by the National Natural Science Foundation of China (No. 32470388; U22A20558), and Shandong Provincial Natural Science Foundation (ZR2024QC197), and fundamental research funds of CAF (Grant No. CAFYBB2023ZA004). Lele Liu is supported by the Cyrus Tang Foundation through the Tang Scholar Program.

## Author contributions

L.Liu. and W.S. planned and designed the research, performed the investigation and data analysis, created visualizations, and wrote the original draft. H.S., C.W., N.L., L.Lin, Y.G., and N.D. contributed to the investigation and manuscript review. W.G. conceived and supervised the research, acquired funding and resources, and reviewed & edited the manuscript. L.Liu. and W.S. contributed equally. All authors critically revised the manuscript and approved its submission.

## Data availability

The complete dataset and associated analysis scripts are permanently hosted on Zenodo under the DOI 10.5281/zenodo.18321077 and can be accessed directly via https://doi.org/10.5281/zenodo.18321077.

## References

1. Alseekh, S., Klemmer, A., Yan, J., Guo, T., & Fernie, A. R. (2025). Embracing plant plasticity or robustness as a means of ensuring food security. Nature Communications, 16(1), 461. 10.1038/s41467-025-55872-4

2. Bates, D., Mächler, M., Bolker, B., & Walker, S. (2015). Fitting linear mixed-effects models using lme4. Journal of Statistical Software, 67(1), 1–48. 10.18637/jss.v067.i01

3. Chen, H., Reed, S. C., Lü, X., Xiao, K., Wang, K., & Li, D. (2021). Coexistence of multiple leaf nutrient resorption strategies in a single ecosystem. Science of The Total Environment, 772, 144951. 10.1016/j.scitotenv.2021.144951

4. Drenovsky, R. E., Pietrasiak, N., & Short, T. H. (2019). Global temporal patterns in plant nutrient resorption plasticity. Global Ecology and Biogeography, 28(6), 728–743. 10.1111/geb.12885

5. Eller, F., Skálová, H., Caplan, J. S., Bhattarai, G. P., Burger, M. K., Cronin, J. T., Guo, W.-Y., Guo, X., Hazelton, E. L. G., Kettenring, K. M., Lambertini, C., McCormick, M. K., Meyerson, L. A., Mozdzer, T. J., Pyšek, P., Sorrell, B. K., Whigham, D. F., & Brix, H. (2017). Cosmopolitan species as models for ecophysiological responses to global change: The common reed *Phragmites australis*. Frontiers in Plant Science, 8, 1833. 10.3389/fpls.2017.01833

6. Ferrer Obiol, J., Bounas, A., Brambilla, M., Lombardo, G., Secomandi, S., Paris, J. R., Iannucci, A., Whiting, J. R., Formenti, G., Bonisoli-Alquati, A., Ficetola, G. F., Galimberti, A., Balacco, J., Batbayar, N., Bragin, A. E., Caprioli, M., Catry, I., Cecere, J. G., Davaasuren, B., Rubolini, D. (2025). Evolutionarily distinct lineages of a migratory bird of prey show divergent responses to climate change. Nature Communications, 16(1), 3503. 10.1038/s41467-025-58617-5

7. Hassani, A., Azapagic, A., & Shokri, N. (2021). Global predictions of primary soil salinization under changing climate in the 21st century. Nature Communications, 12(1), 6663. 10.1038/s41467-021-26907-3

8. Huang, Y., Lack, J. B., Hoppel, G. T., & Pool, J. E. (2022). Gene regulatory evolution in cold-adapted fly populations neutralizes plasticity and may undermine genetic canalization. Genome Biology and Evolution, 14(4), evac050. 10.1093/gbe/evac050

9. Jiang, L., Wang, H., Li, S., Dai, X., Meng, S., Fu, X., Yan, H., Zheng, J., Ma, N., & Kou, L. (2023). A ‘Get-Save-Return’ process continuum runs on phosphorus economy among subtropical tree species. Journal of Ecology, 111(4), 861–874. 10.1111/1365-2745.14066

10. Killingbeck, K. T. (1996). Nutrients in senesced leaves: Keys to the search for potential resorption and resorption proficiency. Ecology, 77(6), 1716–1727. 10.2307/2265777

11. Kobe, R. K., Lepczyk, C. A., & Iyer, M. (2005). Resorption efficiency decreases with increasing green leaf nutrients in a global data set. Ecology, 86(10), 2780–2792. 10.1890/04-1830

12. Lai, J., Zou, Y., Zhang, S., Zhang, X., & Mao, L. (2022). glmm.hp: An R package for computing individual effect of predictors in generalized linear mixed models. Journal of Plant Ecology, 15(6), 1302–1307. 10.1093/jpe/rtac096

13. Li, J., & Prentice, I. C. (2024). Global patterns of plant functional traits and their relationships to climate. Communications Biology, 7(1), 1136. 10.1038/s42003-024-06777-3

14. Liu, L., Pei, C., Liu, S., Guo, X., Du, N., & Guo, W. (2018). Genetic and epigenetic changes during the invasion of a cosmopolitan species (*Phragmites australis*). Ecology and Evolution, 8(13), 6615–6624. 10.1002/ece3.4144

15. Liu, L., Yin, M., Guo, X., Wang, J., Cai, Y., Wang, C., Yu, X., Du, N., Brix, H., Eller, F., Lambertini, C., & Guo, W. (2022). Cryptic lineages and potential introgression in a mixed-ploidy species (*Phragmites australis*) across temperate China. Journal of Systematics and Evolution, 60(2), 398–410. 10.1111/jse.12672

16. Liu, L., Yin, M., Guo, X., Yu, X., Song, H., Eller, F., Ma, X., Liu, X., Du, N., Wang, R., & Guo, W. (2021). The river shapes the genetic diversity of common reed in the Yellow River Delta via hydrochory dispersal and habitat selection. Science of The Total Environment, 764, 144382. 10.1016/j.scitotenv.2020.144382

17. Liu, L., Yin, M., Guo, Y., Song, H., Guo, X., & Guo, W. (2025). Climatic adaptation and phylogenetic history shape the intra-specific variation of CSR strategies in a widespread grass. Plant Diversity. 10.1016/j.pld.2025.06.001

18. Luo, Y., Chen, Y., Peng, Q., Li, K., Mohammat, A., & Han, W. (2021). Nitrogen and phosphorus resorption of desert plants with various degree of propensity to salt in response to drought and saline stress. Ecological Indicators, 125, 107488. 10.1016/j.ecolind.2021.107488

19. Meyerson, L. A., Cronin, J. T., Packer, J., Pyšek, P., & Saltonstall, K. (2025). Ecology, evolution, and North American invasion of one of the world’s most successful plant species. *Annual Review of Ecology*, Evolution, and Systematics. 10.1146/annurev-ecolsys-102723-051221

20. Nicotra, A. B., Atkin, O. K., Bonser, S. P., Davidson, A. M., Finnegan, E. J., Mathesius, U., Poot, P., Purugganan, M. D., Richards, C. L., Valladares, F., & van Kleunen, M. (2010). Plant phenotypic plasticity in a changing climate. Trends in Plant Science, 15(12), 684–692. 10.1016/j.tplants.2010.09.008

21. Prieto, I., & Querejeta, J. I. (2020). Simulated climate change decreases nutrient resorption from senescing leaves. Global Change Biology, 26(3), 1795–1807. 10.1111/gcb.14914

22. Radersma, R., Noble, D. W. A., & Uller, T. (2020). Plasticity leaves a phenotypic signature during local adaptation. Evolution Letters, 4(4), 360–370. 10.1002/evl3.185

23. Ramirez-Parada, T. H., Park, I. W., Record, S., Davis, C. C., Ellison, A. M., & Mazer, S. J. (2024). Plasticity and not adaptation is the primary source of temperature-mediated variation in flowering phenology in North America. Nature Ecology & Evolution, 8(3), 467–476. 10.1038/s41559-023-02304-5

24. Ren, L., Guo, X., Liu, S., Yu, T., Guo, W., Wang, R., Ye, S., Lambertini, C., Brix, H., & Eller, F. (2020). Intraspecific variation in *Phragmites australis*: Clinal adaption of functional traits and phenotypic plasticity vary with latitude of origin. Journal of Ecology, 108(6), 2531–2543. 10.1111/1365-2745.13401

25. Saltonstall, K. (2002). Cryptic invasion by a non-native genotype of the common reed, *Phragmites australis*, into North America. Proceedings of the National Academy of Sciences, 99(4), 2445–2449. 10.1073/pnas.032477999

26. Seibert, S. L., Greskowiak, J., Oude Essink, G. H. P., & Massmann, G. (2024). Understanding climate change and anthropogenic impacts on the salinization of low-lying coastal groundwater systems. Earth’s Future, 12(8), e2024EF004737. 10.1029/2024EF004737

27. Sun, X., Kang, H., Chen, H. Y. H., Björn, B., Samuel, B. F., & Liu, C. (2016). Biogeographic patterns of nutrient resorption from *Quercus variabilis* Blume leaves across China. Plant Biology, 18(3), 505–513. 10.1111/plb.12420

28. Sun, X., Li, D., Lü, X., Fang, Y., Ma, Z., Wang, Z., Chu, C., Li, M., & Chen, H. (2023). Widespread controls of leaf nutrient resorption by nutrient limitation and stoichiometry. Functional Ecology, 37(6), 1653–1662. 10.1111/1365-2435.14318

29. Vahsen, M. L., Blum, M. J., Megonigal, J. P., Emrich, S. J., Holmquist, J. R., Stiller, B., Todd-Brown, K. E. O., & McLachlan, J. S. (2023). Rapid plant trait evolution can alter coastal wetland resilience to sea level rise. Science, 379(6630), 393–398. 10.1126/science.abq0595

30. Vergutz, L., Manzoni, S., Porporato, A., Novais, R. F., & Jackson, R. B. (2012). Global resorption efficiencies and concentrations of carbon and nutrients in leaves of terrestrial plants. Ecological Monographs, 82(2), 205–220. 10.1890/11-0416.1

31. Wakeel, A., Farooq, M., Qadir, M., & Schubert, S. (2011). Potassium substitution by sodium in plants. Critical Reviews in Plant Sciences, 30(4), 401–413. 10.1080/07352689.2011.587728

32. Wang, C., Liu, L., Yin, M., Eller, F., Brix, H., Wang, T., Salojärvi, J., & Guo, W. (2024). Genome-wide analysis tracks the emergence of intraspecific polyploids in *Phragmites australis*. Npj Biodiversity, 3(1), 1–15. 10.1038/s44185-024-00060-8

33. Warton, D. I., Duursma, R. A., Falster, D. S., & Taskinen, S. (2012). Smatr 3– an R package for estimation and inference about allometric lines. Methods in Ecology and Evolution, 3(2), 257–259. 10.1111/j.2041-210X.2011.00153.x

34. Yang, G., Deng, M., Guo, L., Du, E., Zheng, Z., Peng, Y., Zhao, C., Liu, L., & Yang, Y. (2025). Characteristics of leaf nutrient resorption efficiency in Tibetan alpine permafrost ecosystems. Nature Communications, 16(1), 4044. 10.1038/s41467-025-59289-x

35. Yuan, Z. Y., & Chen, H. Y. H. (2015). Negative effects of fertilization on plant nutrient resorption. Ecology, 96(2), 373–380. 10.1890/14-0140.1

36. Zhang, Z., Zheng, J., Guang, Y., Chen, L., Luo, X., Chen, D., & Hu, X. (2022). Phenotypic plasticity contributes more to the variations in nutrient resorption than genetic differentiation in a grassland dominant. Functional Ecology, 36(10), 2605–2615. 10.1111/1365-2435.14152

37. Zhao, K. F., Feng, L. T., & Zhang, S. Q. (1999). Study on the salinity-adaptation physiology in different ecotypes of *Phragmites australis* in the Yellow River Delta of China: Osmotica and their contribution to the osmotic adjustment. Estuarine, Coastal and Shelf Science, 49, 37–42. 10.1016/S0272-7714(99)80006-7

38. Zuo, Z., Reich, P. B., Qiao, X., Zhao, H., Zhang, L., Yang, L., Lv, T., Tang, Z., Yu, D., & Wang, Z. (2024). Coordination between bioelements induce more stable macroelements than microelements in wetland plants. Ecology Letters, 27(11), e70025. 10.1111/ele.70025

