## Supplementary material for "Genetic canalization of nutrient resorption: evidence from a widespread grass under effective salt stress": S

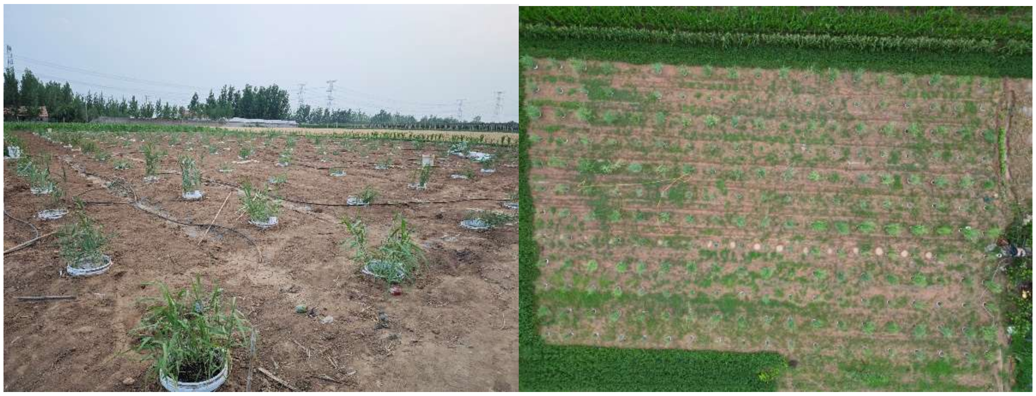

Figure S1 The common garden pictures in a early stage.

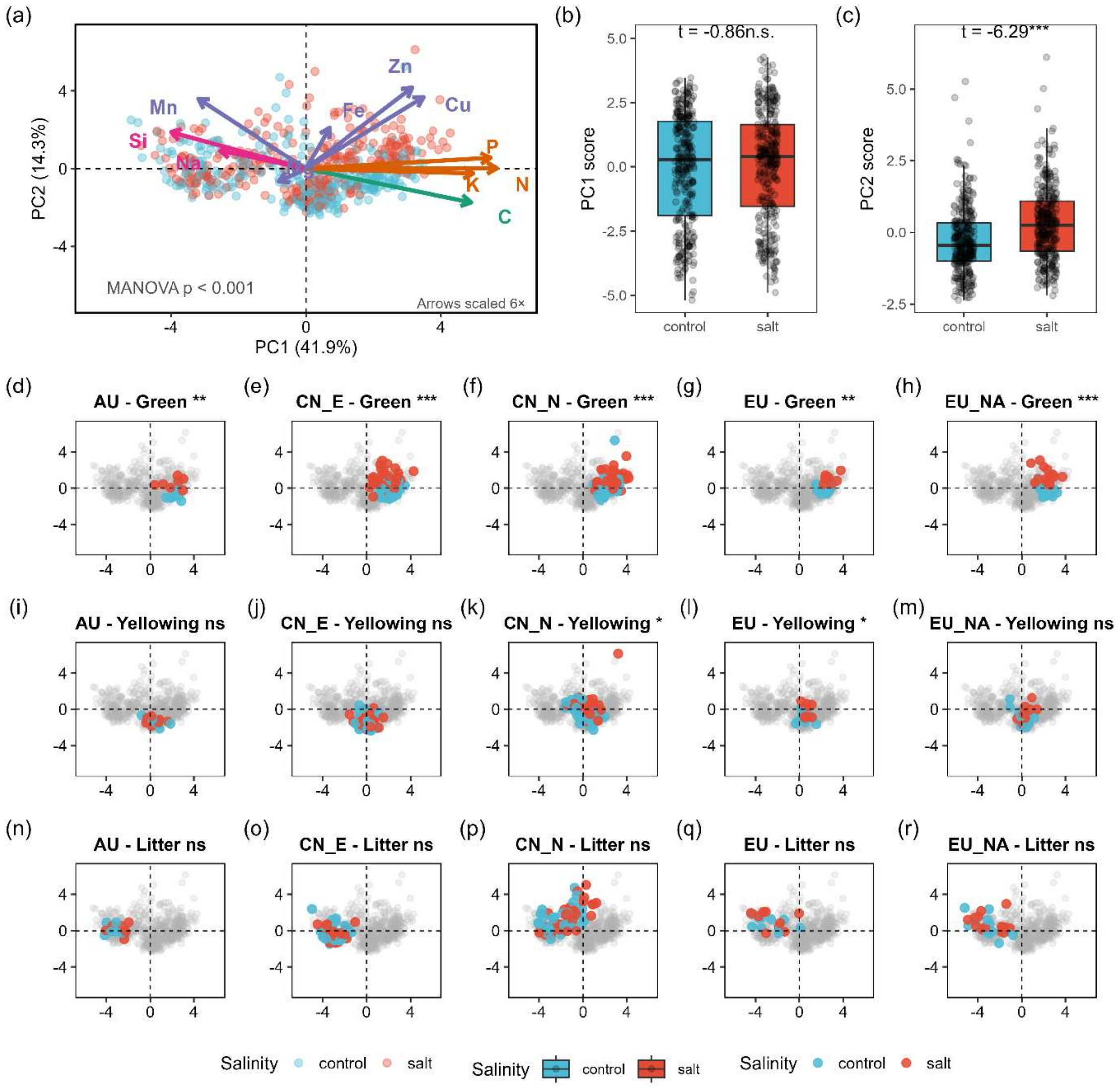


Figure S2 Principal component analysis (PCA) of leaf elemental composition across *Phragmites australis* lineages, salinity treatments, and developmental stages. (a) Global PCA of all samples, with points colored by salinity treatment (control: blue, salt: red). Arrows represent element loadings scaled by a factor of 3; arrow colors indicate functional groups of elements. The MANOVA test (Pillai's trace) revealed a significant overall effect of salinity on PC1 and PC2 combined ($p < 0.001$). (bc) Boxplots of PC1 and PC2 scores comparing control and salt treatments, with t-test results (Welch's t-test) displayed above each panel (* $p$ < 0.05, ** $p$ < 0.01, *** $p$ < 0.001, ns: not significant). (d-f) PCA subplots for each combination of lineage (AU, CN_E, CN_N, EU, EU_NA) and developmental stage (Green, Yellowing, Litter). Background gray points represent all samples, while colored points highlight the focal lineage × stage combination, with colors indicating salinity treatment. The significance of salinity effect within each combination (MANOVA on PC1 and PC2) is shown in the subplot title (*** $p$ < 0.001, ** $p$ < 0.01, * $p$ < 0.05, ns: not significant).

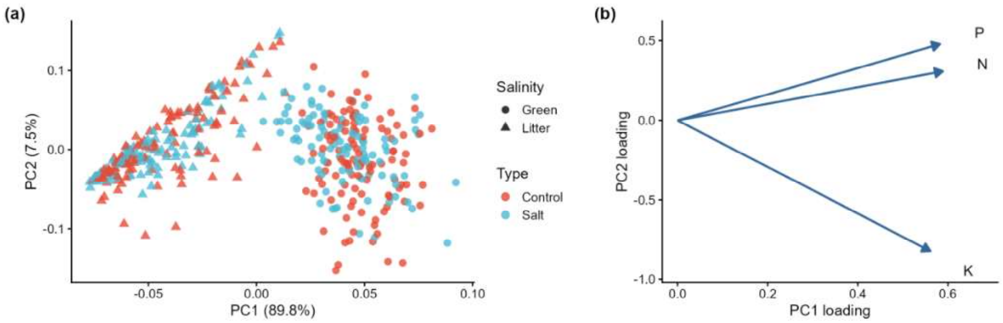


Figure S3 Principal component analysis of nitrogen (N), phosphorus (P), and potassium (K) concentrations in green leaves and leaf litter. (a) Ordination of samples from control and salt treatments across the first two principal components (PC1 and PC2). (b) Vector loadings of individual elements, indicating their contribution and direction of influence on the principal components.

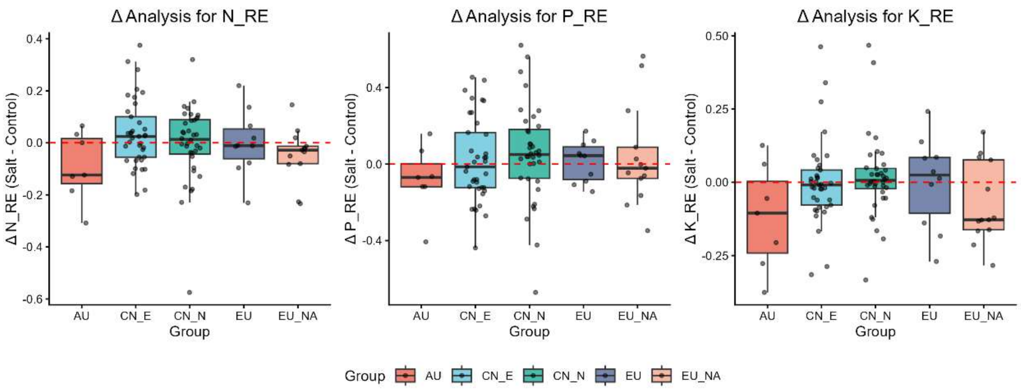


Figure S4 Differences in Nutrient Resorption Efficiency (Δ = Salt – Control) Among Phylogeographic Groups of *Phragmites australis*. Box plots showing the within-genotype difference (Δ) in nitrogen (N), phosphorus (P) and potassium (K) resorption efficiency between salt and control treatments for each phylogeographic group. The Δ value was calculated as the response under salt treatment minus the response under control treatment for the same genotype. The horizontal dashed line at Δ = 0 indicates no change. Boxes represent the interquartile range (IQR), the middle line denotes the median, and whiskers extend to 1.5× IQR. Individual points represent values for each genotype. Positive Δ values indicate higher resorption under salt treatment, whereas negative values indicate lower resorption. No significant differences were found among phylogeographic groups in their Δ values for N, P or K (one-way ANOVA, $p > 0.05$).

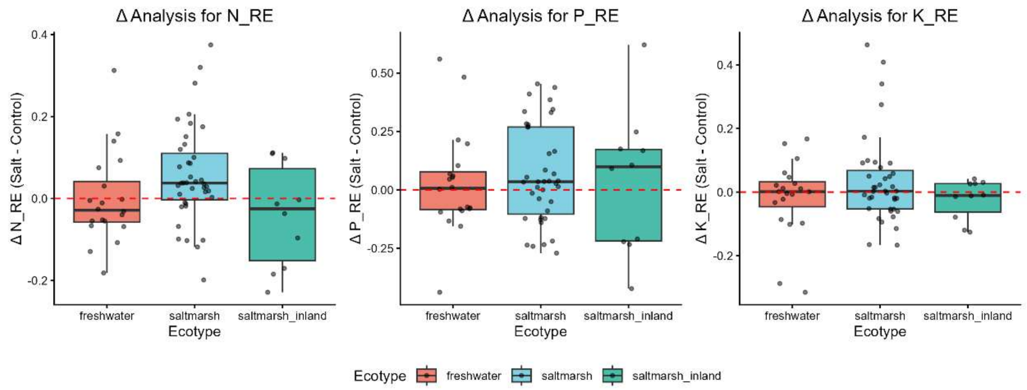


Figure S5 Differences in Nutrient Resorption Efficiency (Δ = Salt – Control) Among Ecotypes of *Phragmites australis*. Box plots showing the within-genotype difference (Δ) in nitrogen (N), phosphorus (P) and potassium (K) resorption efficiency between salt and control treatments for each ecotype. The Δ value was calculated as the response under salt treatment minus the response under control treatment for the same genotype. The horizontal dashed line at Δ = 0 indicates no change. Boxes represent the interquartile range (IQR), the middle line denotes the median, and whiskers extend to 1.5× IQR. Individual points represent values for each genotype. Positive Δ values indicate higher resorption under salt treatment, whereas negative values indicate lower resorption. No significant differences were detected among ecotypes in their Δ values for N, P or K (one-way ANOVA, $p > 0.05$)

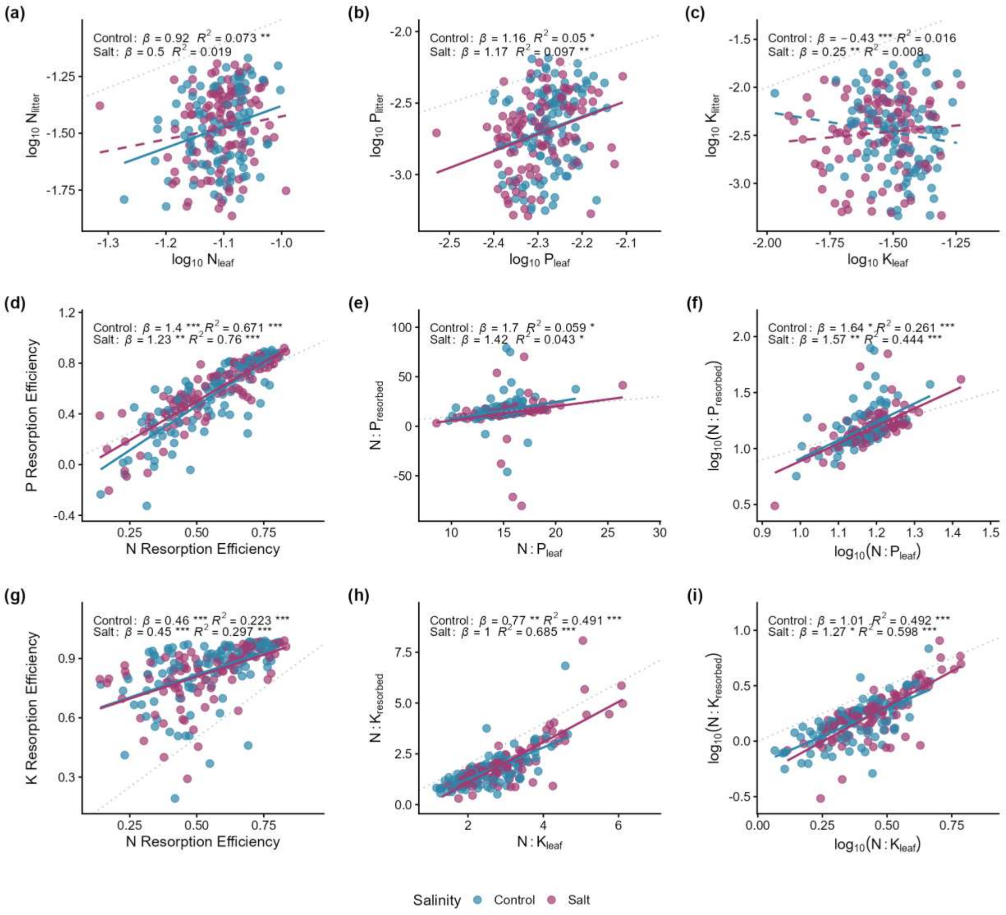


Figure S6 Assessment of three control strategies for nutrient resorption in *Phragmites australis* under varying salinity levels using ordinary least squares (OLS) regression. (a–c) Relationship between nutrient concentrations (N, P, K) in green leaves and senesced litter. (d, g) Association between phosphorus and nitrogen resorption efficiencies. (e, h) Linkage between resorbed N:P(K) ratios and green leaf N:P(K) ratios. (f, i) Correlation between $\log_{10}$-transformed resorbed N:P(K) and $\log_{10}$-transformed green leaf N:P(K) ratios. The gray dashed line in each panel indicates the 1:1 reference. Solid lines represent significant relationships ($p < 0.05$ for the OLS regression model), while dashed lines indicate non-significant relationships. Statistical annotations show results from OLS regression for each salinity level: $\beta$ (with asterisks) is the OLS slope, where asterisks after $\beta$ indicate significant deviation from 1 (* $p < 0.05$, ** $p < 0.01$, *** $p < 0.001$), and asterisks after $R^2$ indicate significance of the OLS regression model.

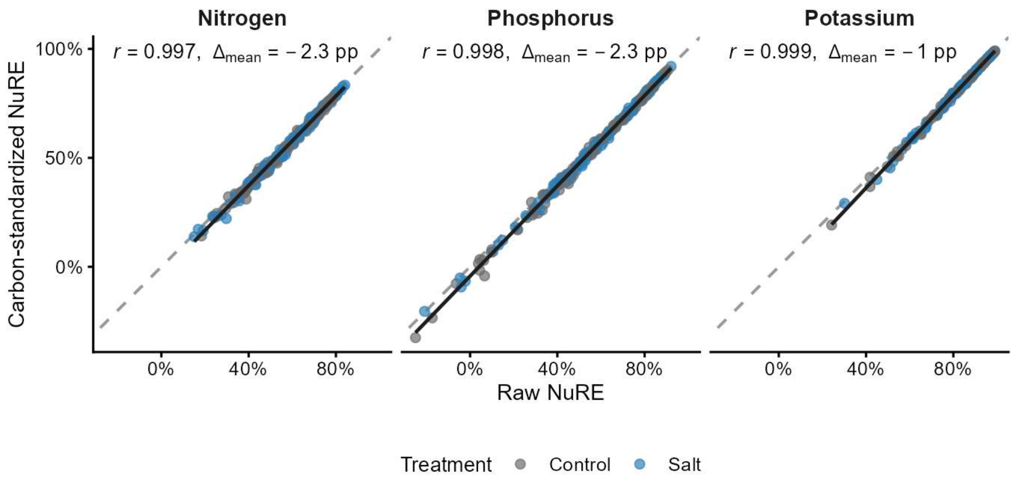


Figure S7 Comparison of raw and carbon-standardized nutrient resorption efficiency (NuRE) for N, P, and K. Points represent genotype-by-treatment observations (*n* = 206), coloured by treatment. Grey dashed lines indicate the 1:1 relationship, and black lines show ordinary least-squares fits. Pearson's *r* and the mean standardized-minus-raw difference (Δmean, percentage points) are shown.

Table S1 The lineages of *Phragmites australis* in the common garden

| Lineage/Ecotype | Lineage description | Sample Size |
|---|---|---|
| AU | Genotypes with haplotype P from Oceania, often named FEAU in the previous papers | 7 |
| CN_E | Genotypes mostly with haplotype P mainly from the Eastern China | 38 |
| CN_N | Genotypes mostly with haplotypes M and O mainly from the Northern China | 37 |
| EU | Genotypes mostly with haplotype M from Europe and Africa | 10 |
| EU_NA | Invasive genotype with haplotype M in the North America | 14 |
| NA | Native genotypes from the North America | 4 |
| Freshwater wetlands | Genotypes originating from non-saline freshwater habitats across China, including the shores of inland rivers and lakes. These environments are characterized by low and stable salinity, relatively nutrient-rich conditions, and seasonal hydrological fluctuations, representing the typical growth conditions for *P. australis* in absence of salt stress. | 20 |
| Coastal saltmarsh | Genotypes collected from tidal salt marshes along the eastern coasts of China. Their habitats are defined by frequent tidal inundation, periodic fluctuations in salinity, and waterlogged, anoxic soils, presenting a dynamic saline stress regime influenced by marine processes. | 40 |
| Inland plateau saltmarsh | Genotypes collected from the hyper-saline environments of the Qaidam Basin on the Qinghai-Tibet Plateau. These habitats are characterized by extreme continental climate, high salinity derived from inland salt lakes, strong UV radiation, and arid conditions, representing one of the most stressful natural habitats for *P. australis* in terms of combined salt and drought stress. | 11 |
